# Resistome and microbiome-immune interactions in an Eastern European population with high antibiotic use

**DOI:** 10.64898/2026.01.22.700835

**Authors:** Bogdan Mirăuță, Anca-Lelia Riza, Ioana Streata, Andrei Pirvu, Stefania Dorobantu, Adina Dragos, Marius Surleac, Mihai G. Netea

## Abstract

The gut microbiome influences host health, affecting gastrointestinal, metabolic, immune, cardiovascular, and neurological functions. A balanced microbiome is associated with favourable health outcomes. However, excessive antibiotic use and dietary habits can disrupt this ecosystem, leading to dysbiosis and affecting body homeostasis.

We present the first comprehensive metagenomic analysis of the gut microbiome in a healthy Romanian cohort. With no prior high-resolution profiling on this population, characterized by high antibiotic consumption, this cohort contributes to understanding microbiome variation in European populations. We report microbiome features consistent with other European populations, including well-defined community configurations, and provide new insights into how these relate to within-phylum diversity. We observe an enrichment of *Enterobacteriaceae*, a pattern that may be shaped by population-level exposures, including antibiotic use.

The analysis of antimicrobial resistance genes, contextualized with data from other cohorts and the European Centre for Disease Prevention and Control, showed an increased prevalence of genes linked to beta-lactams, macrolides, and quinolones—antibiotics commonly used in this population.

Finally, we investigate the relationship between the microbial profile and the systemic immune responses, inferred from correlations with *in vitro* cytokine production. Notably, we identify a potential immune-priming role for *Collinsella* species and a link between the *Prevotella* enterotype and the cytokine production capacity.

**Importance:** This first comprehensive study of the healthy gut microbiome in a Romanian cohort contributes to a baseline for the microbiome and resistome composition of this population. While universally accepted definitions of “healthy” microbiomes, or baseline resistomes, remain lacking, such data help contextualise future studies and support the monitoring of dynamics.

The *Enterobacteriaceae* abundance suggests a microbiome composition potentially influenced by antimicrobial consumption, a relevant pattern in a region with a high burden of nosocomial infections. In addition, the prevalence of antimicrobial resistance genes, and the concordance with commonly used antibiotics in the community reinforces the need to address antibiotic use in public health strategies.

Although links between the gut microbiome and host immunity are not fully understood, our findings are consistent with a role for microbiome composition in immune-related traits. The association between the *Prevotella* enterotype and cytokine balance may provide a basis for further investigation of enterotype-specific immune characteristics.

## Introduction

The human gut microbiome (GM) is now widely recognized as playing an important role in human health. Gut microbiome composition and diversity are influenced by multiple factors, including host genetics, age, lifestyle, diet, and medication use (Lozupone et al. 2012; Heiman and Greenway 2016; Goodrich et al. 2014; Jeffery et al. 2016). Thanks to advances in sequencing technologies and analytical tools, we now have a detailed understanding of GM composition (Almeida et al. 2021). Large-scale gut microbiome cohorts are now relatively common (Aasmets et al. 2022; Gacesa et al. 2022), but they remain concentrated in a few populations (Abdill et al. 2022). This limits our understanding of microbiome variation across ecological contexts and can produce population profiles with gaps, affecting both research and health policies. Previous studies on the human GM in populations from Eastern Europe such as Romania had mainly a focus on understanding disease-associated microbiome alterations (Mihele et al. 2025; Gradisteanu Pircalabioru et al. 2022) or the enteric pathogens (Rodríguez-Molina et al. 2021), but no comprehensive characterisation of the microbiome in healthy cohorts has been performed.

Historically, antimicrobial use in Romania has been reported to exceed the European Union average as reported by ECDC, the European Centre for Disease Prevention and Control, (ECDC 2020b) for many antibiotic groups, and monitoring of antimicrobial resistance genes (ARGs) in key reservoirs has been undertaken in recent years, although primarily focusing on wastewater and environmental samples (Szekeres et al. 2017; Berglund et al. 2023) or on the in depth characterisation of pathogen strains (Popa et al. 2021). No publicly available characterisation of the ARGs in the human gut microbiome currently exists for healthy Romanian donors. The gut microbiome harbours a dense and diverse microbial community, providing a setting for both commensal and opportunistic bacteria and frequent horizontal gene transfer, making it a major reservoir of antibiotic resistance genes that can be mobilized to pathogens (Shoemaker et al. 2001; Garcia-Gutierrez et al. 2019). In addition, antibiotic consumption at population level has been linked to enrichment of ARGs in the human gut (Lee et al. 2023) and a long term lower diversity was reported for individuals after macrolides consumption (Mulder et al. 2020). Characterizing the gut resistome in healthy individuals can reveal baseline ARG distribution, potential dissemination pathways, and the ecological pressures shaping their dynamics.

In addition, the gut microbiome influences systemic immune responses (Schirmer et al. 2016), with several mechanisms being proposed: immunomodulatory effects of microbial metabolites such as short-chain fatty acids (SCFAs), or immune priming effects of certain microbial cell-wall components such as peptidoglycans and lipopolysaccharides (Clarke et al. 2010). Alterations in the gut microbiome (GM) have been increasingly associated in the literature with established immune–mediated diseases. In inflammatory bowel disease, microbial co–abundance networks are disrupted (Geese et al. 2025), and changes in taxa abundance, such as *Ruminococcus*, *Blautia*, *Colidextribacter*, *Oscillospiraceae* and *Roseburia*, have been linked to disease onset (Garay et al. 2023). In rheumatoid arthritis, dysbiosis is associated with Th17–cell–mediated inflammation: elevated *Prevotella copri* has been observed in early RA patients (Maeda and Takeda 2017) and *Collinsella* correlates with increased inflammatory activity (Ruiz-Limón et al. 2022). In multiple sclerosis, patients showed reductions in *Faecalibacterium* and *Prevotella*, alongside an increase in *Actinomyces* (Lin et al. 2024).

The gut microbiome has also been associated with in-vitro PBMC cytokine production in healthy individuals (Schirmer et al. 2016). Dominant patterns of microbiome variation have been proposed to underlie inter-individual differences in cytokine production capacity, with specific taxa and functional profiles correlating with distinct cytokine responses. Although the effect sizes identified to date are modest, these findings support the hypothesis that the gut microbiome may exert long-term immunomodulatory effects on the host immune system, contributing to the shaping of immune responsiveness and susceptibility to immune-mediated diseases.

In this study, we present a comprehensive analysis of the gut microbiome in a healthy Romanian cohort. We describe the overall microbiome composition, assess whether previously defined enterotypes are present in this population and their link with the community structure, and characterize the variability in microbial profiles, including comparisons with other population-level datasets. We further examine the composition and prevalence of the gut resistome. Finally, we explore the relationship between the microbiome and systemic immune responses, and evaluate the cross-population consistency of these associations.

## Results

### Intestinal microbiome of a Romanian population

We sequenced microbiome stool samples from 215 donors out of which after QC (see Methods) we retained 184 samples with more than 30 mil reads for downstream analyses. The mapping to the Unified Human Gastrointestinal Genome (Almeida et al. 2021) resulted in 2915 taxa identified in at least one sample with 525 taxa detected on average in each sample (Supplementary Table 1). This corresponds to 697 bacterial genera detected in at least one sample, 174 genera on average in each sample, 49 genera consistently detected (>50% of samples; >0.1% abundance), out of which 13 corresponded to uncharacterized genera from *Clostridia* class (*Oscillospirales* and *Lachnospirales*). In 28 samples, *Prevotella* (13), *Bacteroides* (10), *Faecalibacterium* (3), *Klebsiella* (1) and *Phocaeicola* (1) genera were dominant (>25%). As expected, *Firmicutes* (*Bacillota*) and *Bacteroidetes* (*Bacteroidota)* are the dominant phyla accounting for approximately 58% and 32% of the bacterial abundance (Figure 1a). The most abundant bacterial genera included *Faecalibacterium*, *Bacteroides*, Prevotella (*Segatella)*, *Phocaeicola* and *Blautia* (Figure 1b).

**Figure 1:**
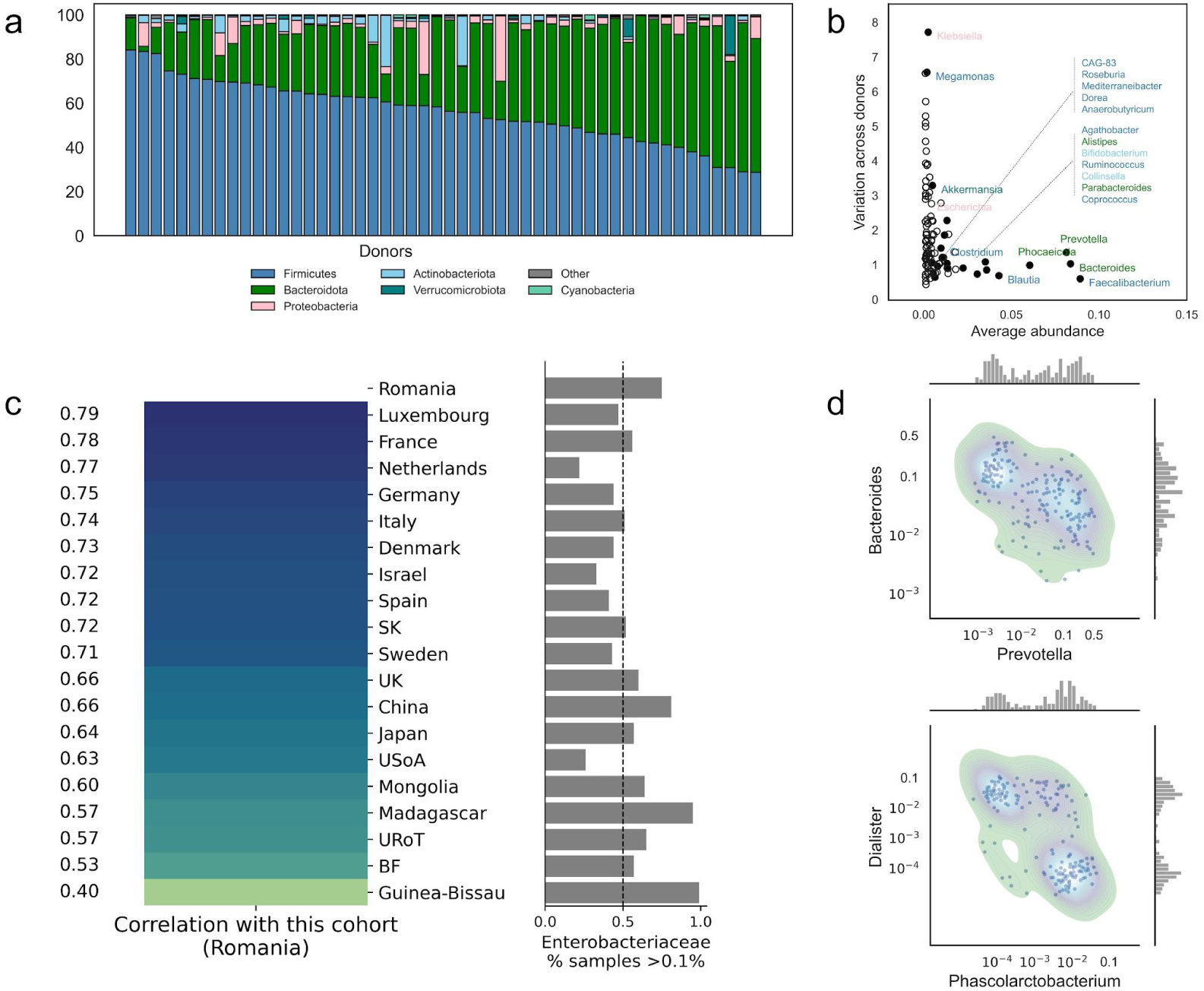
a) Gut microbiome (GM) phylum composition. Samples are ordered by the relative abundance of *Firmicutes* (*Bacillota)*. 50 samples with the highest read coverage are shown. b) Abundance vs. variance of major bacterial genera. The x-axis represents the average abundance of genera (as a percentage of total mapped reads), while the y-axis indicates variability across samples (coefficient of variation; cv). Genera are colored by phylum according to the legend from A. We show genera detected in more than 20 samples and > 0.1% average relative abundance. c) Comparison with external cohorts. Heatmap of bacterial-family–level correlations, showing mean pairwise correlations across individuals. The barplot on the right shows the fraction of samples with more than 0.1% *Enterobacteriaceae.* d) Microbiome composition types. Distribution of log abundance for the genera *Bacteroides* - *Prevotella* and *Dialister* - *Phascolarctobacterium*.

A second mapping to Refseq, the KrakenPlusPF database, (Wood et al. 2019) resulted in mapping 50% of reads to more than 14,000 taxa (Supplementary Table 2). This mapping, extended also to viruses, detected in half of the samples (> 1% in 7 samples) *Carjivirus* and *Burzaovirus*, members of the order *Crassvirales* and among the most prevalent dsDNA bacteriophages of *Bacteroides* in the human gut (Smith et al. 2023).

Romanian gut microbiome composition has a better alignment with European than with non-European samples from the previous studies (Yin et al. 2025) (see Figure 1c). Core taxa (median abundance >1%) were identified at comparable thresholds, both in this cohort and in countries with similar population characteristics (Supplementary Table 3). We observed concordance in genus-level abundances, alongside reduced abundance variation within our cohort (Supplementary Figure 1). This includes major *Firmicutes* genera such as *Faecalibacterium* and *Blautia* (coefficient of variation < 0.7). We note, however, that this lower variation may be the result of this cohort being from a geographically more homogeneous population.

This population exhibits a significant *Prevotella-Bacteroides* dichotomy, genera dominated by *Prevotella sp900557255* (24% of genera abundance) and *Bacteroides uniformis* (30%) respectively. This is consistent with the known enterotypes (Arumugam et al. 2011) and the expected industrial and agrarian nutritional mix in the Romanian population. We also note in this population clear succinotypes (Anthamatten et al. 2024) with individuals having either *Phascolarctobacterium* or *Dialister* (r = – 0.6). While *Dialister* showed multiple significant correlations with other taxa, *Phascolarctobacterium* appeared more ecologically isolated (Supplementary Table 4). Both succinate consumers show increased abundance in the *Prevotella* enterotype, a major succinate producer. The abundance of *Bacteroides* is negatively associated with alpha diversity within *Firmicutes* but positively with diversity within *Bacteroidota*, and conversely for *Prevotella* (Supplementary Figure 2; Methods). This pattern, consistent with the known effects of fiber-rich diets, metabolite production and the more generalist role of *Bacteroides* compared to *Prevotella*, is also observed in other cohorts (Supplementary Table 5). In addition, the *Firmicutes* diversity is associated (r=0.6) with CAG-170 and CAG-177 abundance, supporting recent predictions of the uncultured genus CAG-170 as a biomarker of health (Silva et al. 2025). We note that associations between phylum-level diversity and constituent taxa should be interpreted as indicating biomarker candidates rather than causal effects, even though direct bacterial effects were excluded from diversity calculations (Methods).

The *Enterobacteriaceae* family was detected constantly at higher levels than in other European countries, with 75% of samples showing more than 0.1% relative abundance, compared to 21-56% in other European countries (Figure 1c). We observe this family in very high abundance (>10%) in 6 samples, with *Klebsiella* reaching dominance (27%) in one individual. The *Escherichia* genus accounted for 75% of this family in median. *Enterobacteriaceae* are associated with opportunistic infections, with many species harboring antibiotic resistance genes (ARGs). Confirming previous results (Yin et al. 2025), we observe that in our dataset species from *Enterococcaceae*, *Lactobacillaceae*, *Streptococcaceae* are co-abundant with *Enterobacteriaceae*, while *Lachnospiraceae* and *Oscillospiraceae* are negatively associated (Fisher enrichment test of associated species; pv<0.01 for all families), enforcing the idea of the relationship between these bacteria and the *Enterobacteriaceae* colonization.

### Resistome

Antibiotic resistance genes (ARGs), biocide and metal resistance genes (BMRGs), as well as virulence factors were mapped against all metagenomic assemblies (Methods; Supplementary Table 5). To contextualize our findings, we compared them with previously published population ARG analyses (Lee et al. 2023), with data from the ECDC, and we investigated the genomic context.

Consistent with the widespread presence of Enterobacteriaceae species in this cohort, 55% of identified resistance determinants were assigned to this family, an important reservoir of ARGs. In general, the distribution of ARG families was similar to that observed in other studies (Figure 2a), with tetracycline-resistant ribosomal protection protein and CfxA beta-lactamase gene families being the most prevalent (>95% of samples).

**Figure 2.**
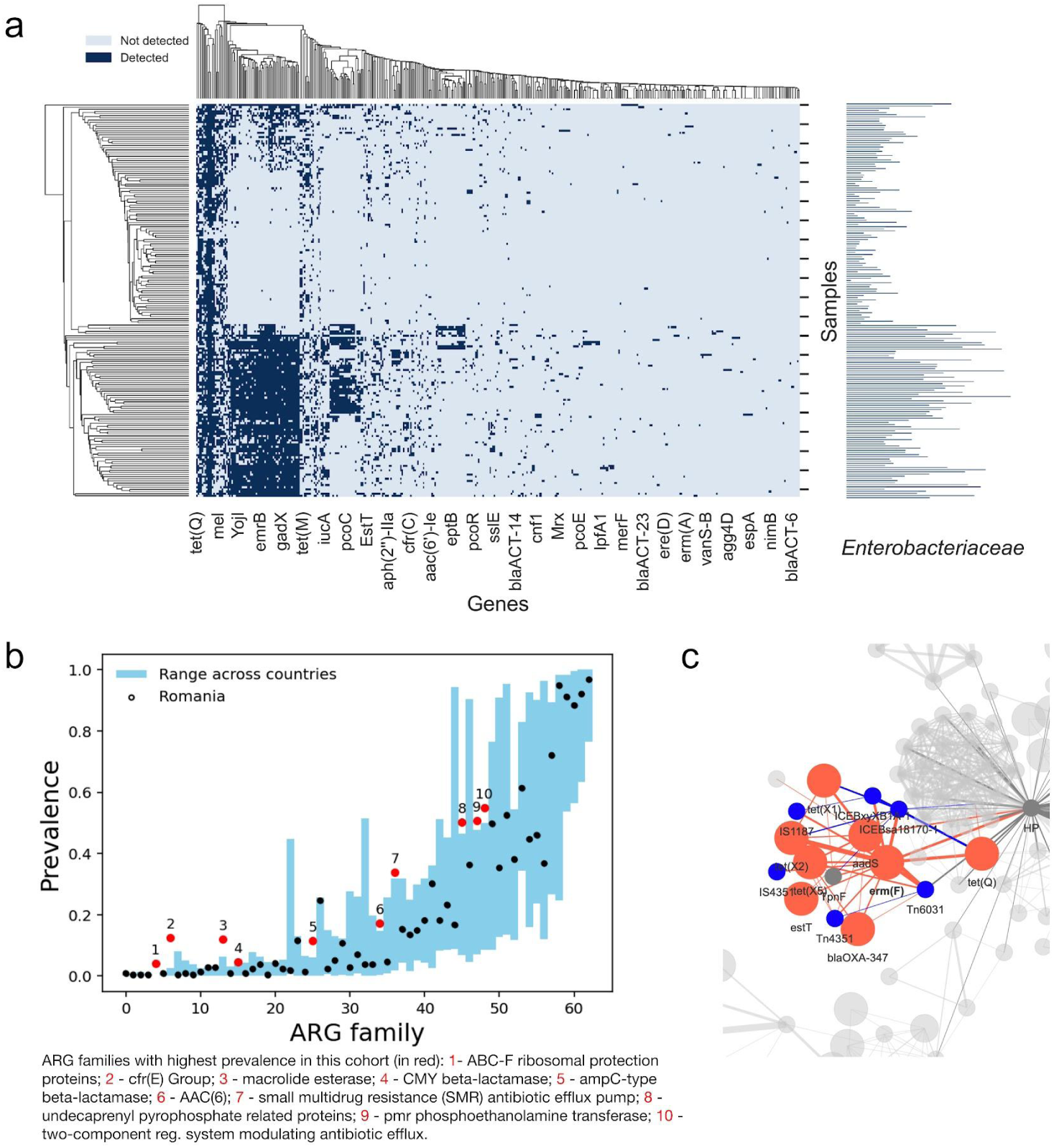
a) Prevalence of antimicrobial resistance gene (ARG) families in this cohort. Left: heatmap of ARG detection in this cohort clustered by genes and samples. An arbitrary selection of gene names is shown on x-axis. Right: *Enterobacteriaceae* family abundance (log) by sample. b) Prevalence of antimicrobial resistance gene (ARG) families in Romania compared with European ranges. For each ARG family (x-axis), the vertical blue bars represent the range of prevalence observed across countries (Lee et al. 2023). Black points indicate the corresponding prevalence values observed in Romania. Red points mark ARG families where Romania’s prevalence is higher than the European range; ARG family annotations is provided under the figure. The y-axis shows prevalence on a scale from 0 to 1. c) Co-localization of resistance determinants. Shown is a snapshot that includes the ARGs (orange) that are associated with *estT*, such as *erm(F)*, *blaOXA-347*, *tet(X5)* and *aadS*, but also various MGEs (blue) such as the Tn6031 transposon like element Tn6031 (blue). Edges indicate genomic proximity (less than <1000 bp apart in the sequence) and the thickness indicates the number of co-occurences. Supplementary Figure 3 expands this view to a genome-wide scale, additionally including biocide resistance genes (yellow) and phage genes (purple).

Out of the 63 gene families assessed, ten showed higher mean prevalence compared to other countries included in this analysis. The contigs for six of these ten families were annotated to *Enterobacteriaceae*, consistent with the observed relatively high prevalence of this family in the cohort. Exceptions included aac(6’), cfr(E) group, and macrolide esterase, which were annotated to *Enterococcaceae*, *Clostridiaceae*, and *Bacteroidaceae* or *Weeksellaceae*, respectively (Blast annotation). No taxonomic assignment could be made for contigs of the miscellaneous ABC-F subfamily ATP-binding cassette ribosomal protection proteins.

We note the high (>0.1) prevalence of the *cfr(E)* gene family, originally characterised in *Clostridioides difficile* isolate DF11 (Stojković et al. 2019). This gene family confers resistance to multiple antibiotics targeting the 23S rRNA, and its detection is consistent with the higher incidence of *C. difficile* infections in hospital settings in this population (Viprey et al. 2023).

The macrolide esterase gene *estT*, a serine-dependent macrolide alpha/beta-hydrolase, was detected in 26 samples, and for two of these individuals we also identified *ereD*, an erythromycin esterase protein. Of note, higher macrolide resistance was also reported by ECDC for *S. pneumoniae* isolates from the Romanian population (ECDC 2020a), despite macrolide in the community consumption being close to the European average (ECDC 2020b). We also note that macrolide resistance potential of this cohort may also be suggested by the high prevalence of ABC-F subfamily genes, which mediate ribosomal protection. While the *eatAv* gene has not been directly linked to macrolide resistance, msr-type ABC-F proteins—though not enriched in our samples—have established associations with macrolide resistance (Ero et al. 2019).

*EstT* was colocalized (on the same contig) with Tn6031, a mobilizable transposon-like element, and, in two samples, in proximity (<1000 bp) of *aadS* or *erm(F)*, also associated with macrolide resistance. We note that, in line with previous observations, *tet(X)* was found associated with Tn6031(Ghosh et al. 2009). Interestingly, in seven samples Tn6031 forms a cluster (pairwise <1000bp in at least one sample) with *aadS, erm(F)*, *tet(X5)* and *blaOXA-347* and IS1187, IS4351 insertion sequences (Figure 2c; Supplementary Figure 3). The colocalization of these genes on the same MGE suggest their co-selection under combined antibiotic pressures and promotes horizontal gene transfer within microbial communities. Other colocalizations involving *bla*_OXA-347_ with macrolide (*erm*(F)) and tetracycline (*tet*(X) variants) resistance genes have previously been associated with poultry production (Liu et al. 2024).

The analysis of antibacterials for systemic use in community (ECDC 2020b) identifies a significantly higher than the EU/EEA mean consumptions in Romania of quinolones and beta lactams. We identify in six individuals (3% of total) the quinolone resistance protein (qnr) family the second highest prevalence among the countries analyzed (Supplementary Table 6).

Confirming the correlation between the beta-lactams consumption and the prevalence of beta-lactamases genes (Lee et al. 2023) we notice high values of most beta-lactamases families in our cohort at similar levels with countries of the high beta-lactam group (France, Italy, Ireland and Spain), with notably an ubiquitous presence of CfxA. The ACI and ampC-type beta-lactamases register the highest difference compared to the European mean with 24% vs 7% and respectively 11% vs 3% prevalence (Supplementary Table 7). The *bla_ACI-1_* gene, an ACI beta-lactamases linked to cephalosporin resistance, appears to be subject to horizontal gene transfer (HGT) via mobile elements (Rands et al. 2018) and was previously identified in plasmids from the rumen gut microbiome (Wu et al. 2025).

The clustering analysis of resistance determinants (Methods; Supplementary Figure 3) identified that betalactamases *blaEC-5/-8/-13/-15* were observed in context with the biocide resistance genes (BRG) such as *sugE*; *blaTEM-7* was observed to cluster with *merP/T/R* mercury resistance genes; *aph(6)-Id*, *dfrA14* and *sul2* resistance genes associate with ICEVchBan9 (an Integrative and Conjugative Element (ICE) from *Vibrio cholerae*). While the ARGs tend to cluster together with other ARGs, MGEs and phage genes, in this case, the BRGs usually cluster together with other BRGs only from the same functional category, such as the *arsA/B/C/D/R*; *soxS/R*; *acrE/S/F*; *mdtO/P/N*.

### Microbiome impact on the systemic cytokine production capacity in peripheral blood mononuclear cells (PBMCs)

Following previous work in Western European populations (Schirmer et al. 2016) (hereon 500FG cohort), we investigated the relationship between the gut microbiome and the systemic host immunity in the present Eastern European cohort. To this purpose, we examined associations between changes in taxa abundances between individuals and the in-vitro cytokine responses of stimulated PBMCs (CS). Cytokine measurements (IL1B, TNFa, IL6, IL1Ra) were performed after PBMC stimulation with LPS, *E. coli*, *C. albicans*, *B. burgdorferi*, *S. aureus*, *M. tuberculosis*, and PHA.

We first applied a linear model to assess the contribution of GM principal components (PCs), the main drivers of variation, to the inter-individual variance in CS levels (Methods). The Ezekiel-corrected coefficients of determination (r²) were within the range previously reported (up to 11%). However, the FDR-adjusted p-values exceeded the significance threshold, indicating that overall microbiome variation is not a predictor of CS changes (Supplementary Table 8).

Next, we analysed the associations between CS and GM abundance at species and genera resolution, and we expanded our analysis to include cytokine profiles averaged by stimulus or cytokine, along with the first 10 cytokine PCs (Supplementary Table 9). We identify five associations at FDR 0.2 (pv=10^-5^) and 84 associations at p<10^-4^.

Interestingly, all taxa except UBA2883, a member of Cyanobacteria, were associated with decreased cytokine responses. Our re-analysis of the 500FG data showed a similar trend, albeit less significant. This pattern supports the hypothesis that the gut microbiome may contribute to immune priming, with observed reductions in cytokine production potentially reflecting that prior host–microbe interactions.

*Collinsella* species were consistently associated with a decreased IL-6 response following bacterial and fungal stimulation. *Collinsella* has been linked to increased inflammation in disease contexts such as rheumatoid arthritis (Ruiz-Limón et al. 2022). The reduced cytokine production in vitro may reflect a distinct, context-dependent function. In a disease-free state *Collinsella* may contribute to fine-tuning the PBMC inflammatory response, supporting immune homeostasis rather than promoting inflammation. Of note, *Collinsella aerofaciens was* identified in the 500FG cohort but was not associated with IL6 production.

*Prevotella* species were associated with Cytokine PC4 (explaining 6% of variance), which correlated positively with IL1B and negatively with TNF-α production capacity, suggesting that *Prevotella*-dominated enterotypes may influence the IL1B–TNF-α balance. Although airway *Prevotella* has been implicated in immune training (Horn et al. 2022), we note that the *Prevotella*-dominant enterotype strongly reflects the host’s diet, which itself may partly explain its association with systemic immunity. A similar negative correlation with the IL1B/TNF-α ratio was observed in the 500FG cohort for *Prevotella copri*, albeit at a lower significance (Supplementary Table 10).

*Bilophila* sp902373525 negatively correlated with Cytokine PC9. The five *Bilophila* species detected, including *B. wadsworthia*, formed a tight cluster (average inter-species *r* = 0.79) and were consistently associated with higher PC9 and lower IL1B and TNF-α production capacity (*p* < 0.05). *B. wadsworthia* has previously been shown to activate innate immunity via TLR2 signaling (Yoshida et al. 2025).

A detailed cross-study replication between the Dutch and Romanian cohorts was limited by differences in analysis pipelines, microbiome annotation and number of taxa identified. We note that we considered 513 species for the 500FG study compared to 4630 taxa included in the present Romanian cohort, resulting in 122 species corresponding to 2,318 associations (Supplementary Table 11). As noted previously, both cohorts showed an enrichment of negative associations between microbial species and cytokine production capacity, with 39 associations being nominally significant (p<0.1) in both studies, and 64% of these showing a consistent effect direction (Fishers exact test, p=0.05).

## Discussion

Our data offers a snapshot into the gut microbiome of healthy individuals from an Eastern European cohort, improving understanding of microbiome variation across Europe and offering insight into a population with historically high antimicrobial drug consumption.

Comparisons with other cohorts show that our data aligns with gut microbiome profiles observed in other European populations. Although geographically confined, this cohort exhibits in nearly equal proportions the *Prevotella* and *Bacteroides* enterotypes. Adding to the extensive literature on diet-microbiome interactions, we find that these enterotypes associate differentially with phylum-level diversity: *Prevotella* correlates with *Firmicutes* diversity, while *Bacteroides* aligns with *Bacteroidota* diversity, patterns that are consistent across other cohorts.

There are, however, some particularities in the Romanian population that emerge from contextualizing our data through comparison with external datasets. In this context, we recognize that both protocol- and analysis-related differences may influence cross-study comparisons. Nevertheless, our use of higher-level resolutions, genius or family, mitigates the impact of methodological variations. Furthermore, the antimicrobial resistance gene analyses are supported by independent taxonomic annotation of selected ARGs using BLAST on contigs, as well as by concordant external evidence from the ECDC.

In this respect, we notice a consistent higher abundance of *Enterobacteriaceae* which likely reflects the historical antibiotic usage in this population or environmental acquisition and may indicate a gut microbiome more prone to perturbation and opportunistic colonization.

The gut microbiome ARG profile from this Romanian cohort, compared to ARG data for European populations, provides complementary insights for national and European ARG monitoring efforts. This key reservoir reflects historical high antibiotic consumption in the community and is a stable component of population-level resistance dynamics. Within this context, we observe signals associated with increased potential resistance to quinolones, beta-lactams, and macrolides, each supported by overrepresented ARG families in our cohort and consistent with patterns reported by ECDC surveillance data. In particular, the enrichment of beta-lactamases and quinolone resistance genes aligns with reports of antimicrobial consumption in the community—a type of use that is likely to exert a stronger influence on ARG prevalence at the population level than hospital-based antibiotic use. This reinforces recent legislative efforts aimed at tightening community antibiotic use, suggesting that such measures are well-targeted and necessary.

We also note that the elevated resistance potential of *C. difficile* may be connected to both overall high antibiotic use, but also the high burden of CDI in hospital settings, from where spores can be disseminated into the wider community. This is in line with our recent survey of the cause of severe infections in tertiary hospitals from the same area of South-West Romania, in which *C. dificille* was a much more frequent cause of sepsis compared to other European countries (Grigorescu et al. 2025). As previously reported for this population (Popa et al. 2021), multidrug-resistant clones have been shown to pass from hospitals into wastewater, even after treatment. These observations highlight the need for stricter control measures.

We report the co–localisation of resistance determinants, which may reflect evolutionary pressures promoting multidrug resistance. In some cases, such as the co–occurrence of *aadS*, *erm*(F), *tet*(X5), and *bla*OXA–347, this pattern suggests potential transmission from agricultural sources, specifically poultry. Existing studies on poultry AMR in Romania, eg. (Brătfelan et al. 2023), are limited in scope, and comprehensive metagenomic datasets are lacking, restricting our ability to determine the role of these mechanisms. Our results underscore the need for such research.

Despite the relatively small cohort size, our study provides additional support to the hypothesis that the gut microbiome can influence immune function. While we did not observe a contribution of global microbiome variation axes to cytokine production capacity in PBMCs, we detected a potential effect on immune responses of the *Prevotella*-associated enterotype. At the genus or species level, we observe limited overlap with previously reported associations, although the consistent enrichment of negative correlations between bacterial abundance and cytokine production capacity suggests a modulating effect of the microbiome on inflammatory responses in both Romanian and Dutch cohorts. This is likely to have an important effect on local homeostasis of the gut by preventing inappropriate induction of local inflammation. Some differences between our results and those of prior studies may reflect variations in both microbiome composition and immune response profiles across populations. As this analysis focused on long-lasting effects, the expected associations are modest. Given the population-level diversity of the gut microbiome (approximately 4,000 species), a systematic assessment would require much larger cohorts, potentially over 1,000 individuals based on power analysis, underscoring the need for more comprehensive studies in larger cohorts.

This dataset covers a geographically limited region, and country-wide data are needed to establish a more systematic and comprehensive map of the gut microbiome and human gut resistome in this population. Data-federation frameworks, such as those within the European life-science infrastructure (Crosswell and Thornton 2012) and the emerging national efforts of the Romanian Bioinformatics Cluster (Mirăuță et al. 2024), could enable broader use of decentralized gut microbiome data for research, complementing health-system monitoring of resistome dynamics.

## Data availability

All referenced tables can be accessed at doi.org/10.5281/zenodo.18324942. Metagenomic assembled contigs corresponding to antimicrobial resistance genes can be accessed at doi.org/10.5281/zenodo.18324942. Microbiome sequencing data are available under ENA project ID PRJEB106858. The cytokine data can be requested from the corresponding author. Access to all datasets will be granted to editors and reviewers during peer review and made publicly available after completion of the peer review process.

## Author Contributions

BM conceptualized the study, performed data analysis, interpreted the results, and wrote the manuscript. MS and BM conducted the ARG data analysis and wrote the corresponding section. ALR, AP, SD, AD, IS collected microbiome samples, and performed cytokine measurements. MN supervised the project and provided revisions. MN and ALR secured funding for all biological data generation. All authors approved the final manuscript.

## Acknowledgement

The analyses of the biological data generated for this article were funded and in the context of “Slowing Immune system Aging through Dietary control” (ImmunAgeD) PNRR/2022/C9/MCID/I8. contract no. 760057/23.05.2023. Raw data was generated through the “Functional Genomic in Severe Sepsis” (FUSE; P_37_745, MySmis 103454, contract no 31/01.09.2016) https://old.umfcv.ro/fuse (accessed on 25 March 2025). The authors would like to thank all patients from the FUSE cohort for their participation in this study. The FUSE study: Netea, M.G.; Ioana, M.; Riza (Costache), A.L.; Dumitrescu, F.; Pîrvu, A.; Streață, I.; Roskanovic, M.; Dorobanțu, S.; Dragoș, A.; Drodar, M.; Cucu, M.-G.; Pleșea, R.-M.; Catalin, B.; Balseanu, A.; Cimpoeru, A; Dragoi, I; Grigorescu, A; Fratea, A.

We acknowledge Alexandru Mizeranschi for providing advice on implementing metagenomic analysis Nextflow pipelines.

We acknowledge the use of Generative Artificial Intelligence (AI) tools, specifically Le Chat (Mistral AI), ChatGPT 4.0 (OpenAI) and Gemini 1.0 (Google) for improving the clarity, grammar, and readability of the manuscript text. No analyses or interpretations were performed using AI, and no conclusions or scientific findings were formulated using AI.

## Ethics approvals

The FUSE study protocol, in which the biological data was generated, was approved by the Committee of Ethics and Academic and Scientific Deontology of the University of Medicine and Pharmacy of Craiova under number 80/17 November 2016 and by the Ethics Committee of “Victor Babeş” Clinical Hospital of Infectious Diseases and Pneumology and performed in accordance with the latest version of the Declaration of Helsinki and guidelines for good clinical practice (GCP).

Informed consent was obtained from all participants following a detailed explanation of the procedures, and potential risks. Participants were provided with information on data usage, confidentiality measures, and their rights as participants. After time for consideration, all participants voluntarily signed a standardized consent form. The signed consent forms are securely maintained at the Human Genomics Laboratory, University of Medicine and Pharmacy of Craiova, Romania.

## Methods

### Cohort

Healthy adult individuals were recruited following informed consent between May 2017 and November 2019 at the Human Genomics Laboratory, University of Medicine and Pharmacy of Craiova. We considered healthy individuals with negative medical history, under no prescribed or self-administered medication. All participants had Eastern European ancestry, and are drawn from a mix of urban (Craiova) and village populations surrounding the city. Biological samples were collected for biobanking (serum/plasma from venous blood, urine and stool samples) or ex-vivo immune experiments (PBMC isolation, stimulation followed by ELISA measurements).

#### Production capacity of proinflammatory cytokines

To capture cytokine responses of immune cells, we measured the production of 24-hr monocyte-derived (IL-1β, TNF-α, IL-6) after stimulation of peripheral blood mononuclear cells [PBMCs] with 8 stimuli (LPS, PHA, heat-killed *Candida albicans, Streptococcus pneumoniae, Escherichia coli, Borrelia burgdorferi, Mycobacterium tuberculosis*).

Ficoll gradient isolation of PBMCs was performed as previously described (Horst et al. 2016). Cells were washed twice in saline and suspended in medium (RPMI 1640) supplemented with gentamicin 10 mg/mL, L-glutamine 10 mM and pyruvate 10 mM. PBMC stimulations were performed with 5×105 cells/well in round-bottom 96-wells plates (Corning) for 24 hr at 37°C and 5% CO2. Additional details are available in the STAR Methods. Supernatants were collected and stored in −20°C until used for ELISA.

ELISA measurements were done using R&D Duoset kits, as per the manufacturers instructions. 450nm and 540nm measurements were done on BMG Clariostar.

### Metagenomics analyses

Paired end (2 x 100 bp) metagenomic sequencing was performed by BGI on the DNBseq platform resulting in 5-6 G (2 x2.5-3) per sample.

Contig assembly was performed using MetaSpades (Nurk et al. 2017) within the nf-core/mag pipeline (Krakau et al. 2022). Following host removal, we performed the taxonomic profiling using Kraken2 (Wood et al. 2019) within the *taxprofiler* nextflow pipeline (Stamouli et al. 2023).

We mapped the taxonomic lineages using GTDB (release202) (Parks et al. 2018). Of note, we used the Prevoletacea family for Prevotella, Alloprevotella Paraprevotella and Hallella genera (Bacteroidaceae in GTDB taxonomy). The RefSeq alignment was performed using K the KrakenPlusPF database downloaded in September 2024 from https://benlangmead.github.io/aws-indexes/k2.

Taxonomic assignments of assembled contigs was performed with Diamond (Buchfink et al. 2015) using the following parameters: diamond blastx -d refseq_bacteria -q [assembled contigs] -o [output file] -b 0.61 --outfmt 6 qseqid sseqid pident length mismatch staxids. For selected contigs we annotated taxa using Blast (megablast) on the nucleotide sequences (McGinnis and Madden 2004).

#### Quality control

For ARG mapping, assembled contigs from all 215 samples were included. All other analyses were restricted to the 184 samples with sequencing depth greater than 30 million reads (median 41 million reads per sample). Association analyses were conducted after removing outlier samples identified by principal component analysis (PCA). PCA was performed on scaled microbiome and cytokine profiles, and outliers were defined as samples exceeding four standard deviations on any principal component. Principal components used in downstream association analyses were recomputed after outlier removal.

#### Phylogenetic entropy

Phylogenetic entropy (Allen et al. 2009) was computed at genera level after removing the taxa whose association with diversity was being tested, and by restricting the calculation to the remaining taxa of interest. Genera abundances were normalized to sum to 1 within each phylum. We used the GTDB bacterial phylogenetic trees (bac120_r226) pruned at genus level. We followed a majority voting approach to map GTDB nodes to the taxa names used in this study).

#### Explained variance

We ran a linear model with the first 20 PCs and calculated the Ezekiel r2 estimator, 1-(N-1)/(N-p-1)x(1-r^2^), where N is the number of samples and p the number of explanatory variables (PCs in this case). P-values were computed using a permutation strategy keeping the PCs unchanged and permuting the cytokine profiles. We then followed a Benjamini Hochberg strategy to account for multiple testing.

#### ARG Methods

The metagenomics assemblies (contigs) have been subject of antibiotic resistance genes (ARG) predictions using CARD (Alcock et al. 2020), ResFinder (Bortolaia et al. 2020) and BacAnt (Hua et al. 2021). Diamond software was used to compare the contig sequences against the protein databases by BlastX (the following command was used: diamond blastx --max-target-seqs 0 --more-sensitive --id 70 -p 8 --subject-cover 90) (Buchfink et al. 2015). The resulting set has at least 70% identity and 90% coverage on the reference sequences. The analysis of the presented results was performed only on the subset having at least 90% to 100% identity. R was used for graph representation. The genomic context clustering has been analyzed with an in house script that looks at genes that are in close proximity in the genome (less than 1000 bp) or in the contig the predicted genes have been found into. The context clustering graph has been generated in R based on the subsets that contained at least two antibiotic or biocide resistance genes in proximity with one another.

Between countries analysis was performed at ARG class resolution. We mapped ARG (CARD ARO Ids) to ARG class by retrieving the parent term in the CARD ontology 4.0.1 (aro.json and aro.obo files). We use prevalence (detection) of ARG or ARG class and average the prevalence across the samples in a country.

## Supplementary Materials

**Supplementary Table 1 - Taxa_Abundance.** CSV file containing taxonomic abundance data. Values are scaled so that abundances within each sample are equal (10 mil reads). Taxonomic lineage information is included for each entry.

**Supplementary Table 2 - Taxa_Abundance_KrakenPlusPF.** CSV file containing taxonomic abundance data following mapping to the KrakenPlusPF database. We report unnormalized read counts for 195 entries in which no dominant genus (>50% relative abundance) was identified. “Root” denotes unaligned reads or reads assigned only to higher-level taxonomic ranks.

**Supplementary Table 3 - Taxonomic_profile_across_cohorts**. CSV file containing taxonomic abundance data for this cohort and cohorts reported in Yin et al., 2025. Values represent relative abundances expressed as percentages, summing to 100% within each sample.

**Supplementary Table 4 - Correlation between taxa**. CSV file reporting the Spearman correlation coefficients between all taxa. Correlations are calculated across all samples in the cohort.

**Supplementary Table 5 - Phylogenetic diversity.** This Excel file contains three sheets: sample_Hp_abund, Bacteroidota_association_bacteria_by_country, and Firmicutes_association_bacteria_by_country. The sample_Hp_abund sheet reports, for each sample and country, the abundances of the top 100 genera, as well as *Firmicutes* and *Bacteroidota* diversity calculated after optional removal of taxa of interest and rescaling. The [Bacteroidota,Firmicutes]_association_bacteria_by_country sheets report correlations between bacterial genera and phylum-level phylogenetic diversity, averaged by country.

**Supplementary Table 6 - ARG** Excel file containing sheets Arg_prevalence, Arg_proximity, Arg_taxa, Contig_arg. Arg_prevalence contains ARG names, additional annotations provided by CARD, and a detection indicator (0 = absent, 1 = present) for each sample in the cohort. Arg_proximity lists pairs of ARGs that are located within 1,000 bp of each other on at least one contig from this cohort. Arg_taxa reports the taxonomic assignments obtained through DIAMOND mapping of taxa to contigs. Contig_arg provides the mapping between contigs and their associated antimicrobial resistance genes (ARGs).

**Supplementary Table 7 - ARG compare Europe**. Excel file containing sheets Aro_arg_family and Arg_family_across_countries. Aro_arg_family maps ARG ARO terms to ARG families Mapping of ARG to ARG families was performed using CARD. Arg_family_across_countries includes the ARG families detected in this study in and in (Lee et al. 2023).

**Supplementary Table 8 - Microbiome PC_cytokine_variance_explained.** Csv file including the variance explained by the first 20 mcirobiome PCs (r2), the probability computed as detailed in Methods (pv), the number of samples on which the association was performed (N) and the fdr corrected value (fdr_bh).

**Supplementary Table 9 – Microbiome_cytokines.** CSV file containing associations between microbiome features and cytokine abundances. Association coefficients (cor) were calculated using Spearman’s rank correlation. The table also includes PC[x]mic and PC[x]cit, representing the principal components of the microbiome and cytokine datasets, respectively. Both Bonferroni- and FDR-adjusted p-values are provided.

**Supplementary Table 10 – Microbiome_cytokines_ro_nld** and **Supplementary Table 11 – Microbiome_cytokines_ro_nld.** These tables contain data analogous to Supplementary Table 9. Supplementary Table 10 includes data reprocessed from the FG500 dataset. Supplementary Table 11 reports associations for microbial species identified in both studies.

**Supplementary Figure 1.**
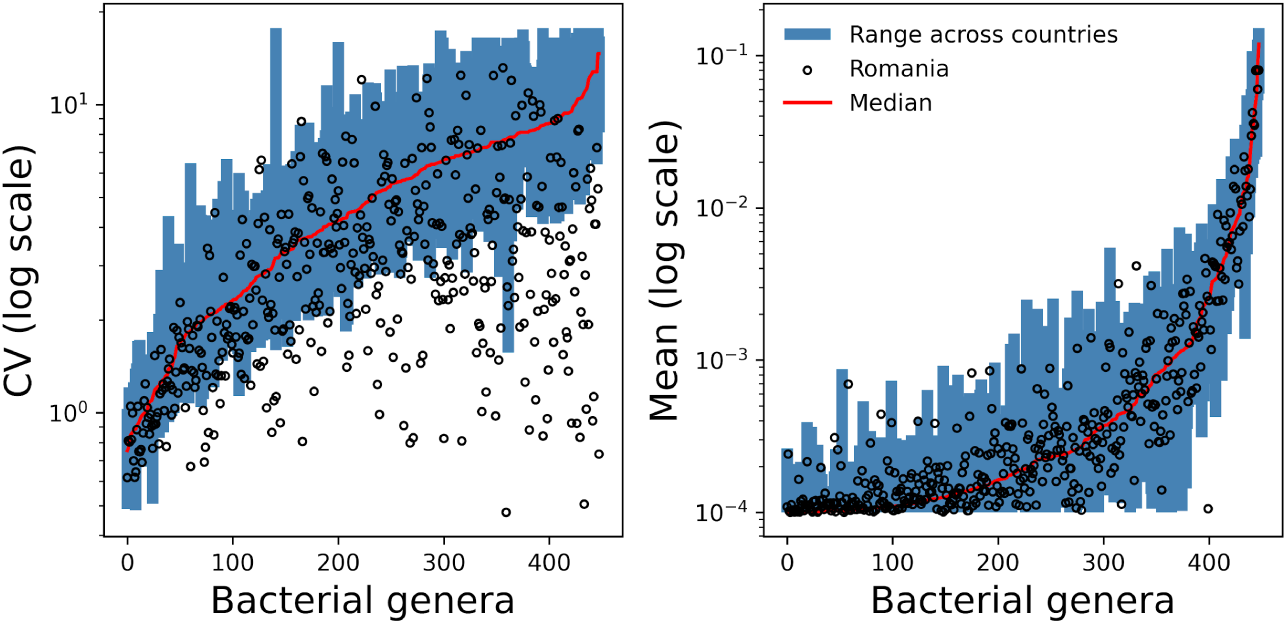
Comparison of taxonomic abundance and variation across cohorts. Mean abundances and variability of bacterial genera (x-axis) in this cohort are compared with those reported in Yin et al., 2025. Blue ranges indicate values observed across the published cohorts, while black circles represent values observed in this cohort. Values are shown on the log scale.

**Supplementary Figure 2.**
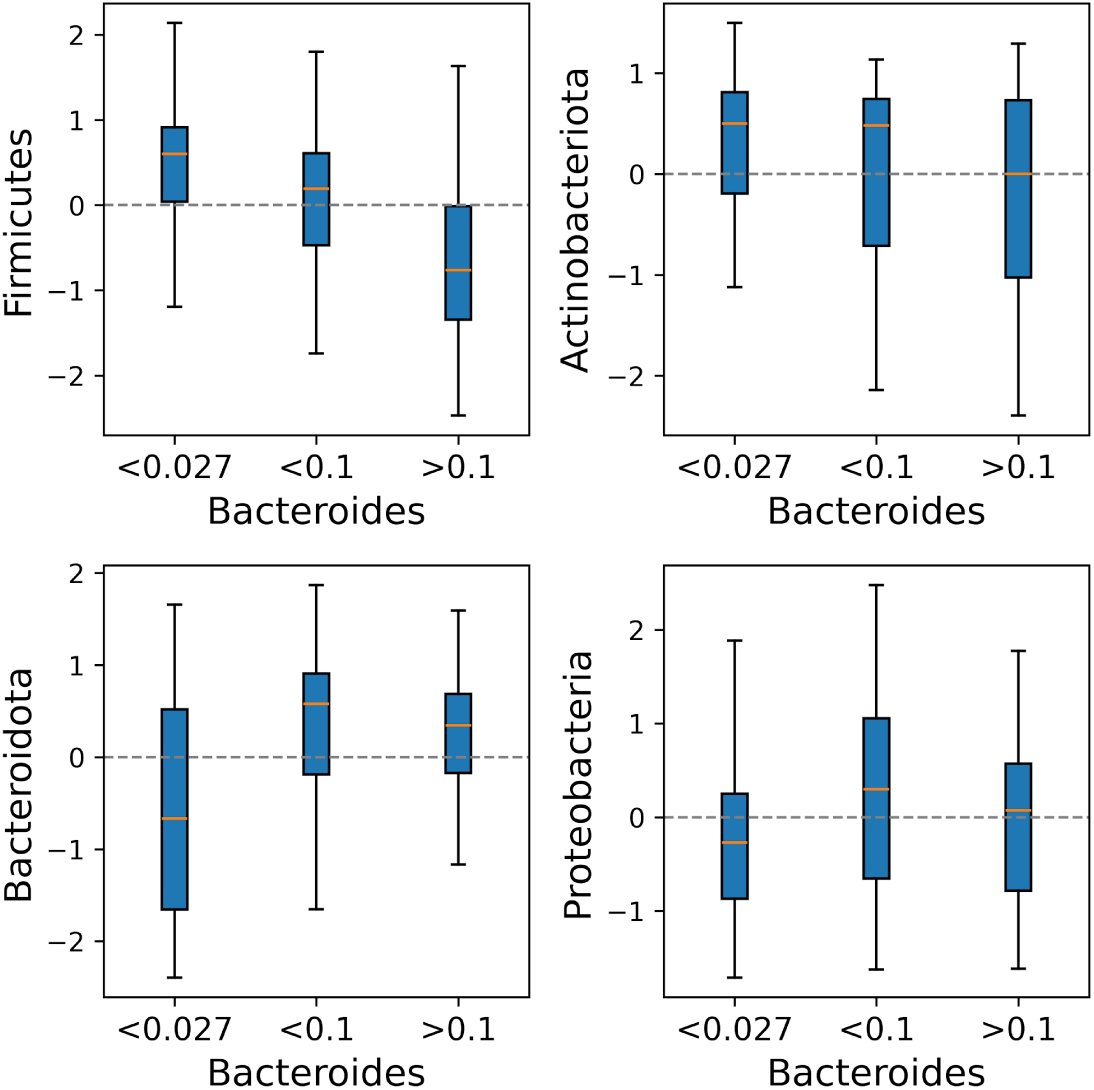
Association between *Bacteroides* abundance and gut microbiome diversity. The y-axis shows phylogenetic entropy (z-scores) calculated for the entire community as well as for taxa belonging to individual bacterial phyla. *Bacteroides* was removed prior to computing community-level diversity, and phylogenetic entropy for each phylum was calculated on corresponding taxa. **An interactive HTML version of this figure is available at** https://zenodo.org/records/1832494

**Supplementary Figure 3.** ARG BMRG MGE context. Co-localization of resistance determinants (Supplement to Figure 2c). Edges represent genomic proximity (<1000 bp apart on a contig), and edge thickness reflects the number of observed co-occurrences in the same contigs. Colors indicate the type of genetic element: orange – ARGs, blue – MGEs, yellow – biocide resistance genes, purple – phage-associated genes. The supplementary HTML file provides the full interactive visualization.

## References

1. Aasmets, Oliver, Kertu Liis Krigul, Kreete Lüll, Andres Metspalu, and Elin Org. 2022. ‘Gut Metagenome Associations with Extensive Digital Health Data in a Volunteer-Based Estonian Microbiome Cohort’. Nature Communications 13 (1): 869. 10.1038/s41467-022-28464-9.

2. Abdill, Richard J., Elizabeth M. Adamowicz, and Ran Blekhman. 2022. ‘Public Human Microbiome Data Are Dominated by Highly Developed Countries’. PLoS Biology 20 (2): e3001536. 10.1371/journal.pbio.3001536.

3. Alcock, Brian P, Amogelang R Raphenya, Tammy T Y Lau, et al. 2020. ‘CARD 2020: Antibiotic Resistome Surveillance with the Comprehensive Antibiotic Resistance Database’. Nucleic Acids Research 48 (D1): D517–25. 10.1093/nar/gkz935.

4. Allen, Benjamin, Mark Kon, and Yaneer Bar-Yam. 2009. ‘A New Phylogenetic Diversity Measure Generalizing the Shannon Index and Its Application to Phyllostomid Bats.’ The American Naturalist 174 (2): 236–43. 10.1086/600101.

5. Almeida, Alexandre, Stephen Nayfach, Miguel Boland, et al. 2021. ‘A Unified Catalog of 204,938 Reference Genomes from the Human Gut Microbiome’. Nature Biotechnology 39 (1): 105–14. 10.1038/s41587-020-0603-3.

6. Anthamatten, Laura, Philipp Rogalla von Bieberstein, Carmen Menzi, et al. 2024. ‘Stratification of Human Gut Microbiomes by Succinotype Is Associated with Inflammatory Bowel Disease Status’. Microbiome 12 (1): 186. 10.1186/s40168-024-01897-8.

7. Arumugam, Manimozhiyan, Jeroen Raes, Eric Pelletier, et al. 2011. ‘Enterotypes of the Human Gut Microbiome’. Nature 473 (7346): 174–80. 10.1038/nature09944.

8. Berglund, Fanny, Daloha Rodríguez-Molina, Gratiela Gradisteanu Pircalabioru, et al. 2023. ‘The Resistome and Microbiome of Wastewater Treatment Plant Workers – The AWARE Study’. Environment International 180 (October): 108242. 10.1016/j.envint.2023.108242.

9. Bortolaia, Valeria, Rolf S. Kaas, Etienne Ruppe, et al. 2020. ‘ResFinder 4.0 for Predictions of Phenotypes from Genotypes’. The Journal of Antimicrobial Chemotherapy 75 (12): 3491–500. 10.1093/jac/dkaa345.

10. Brătfelan, Dariana Olivia, Alexandra Tabaran, Liora Colobatiu, Romolica Mihaiu, and Marian Mihaiu. 2023. ‘Prevalence and Antimicrobial Resistance of Escherichia Coli Isolates from Chicken Meat in Romania’. Animals: An Open Access Journal from MDPI 13 (22): 3488. 10.3390/ani13223488.

11. Buchfink, Benjamin, Chao Xie, and Daniel H. Huson. 2015. ‘Fast and Sensitive Protein Alignment Using DIAMOND’. Nature Methods 12 (1): 59–60. 10.1038/nmeth.3176.

12. Clarke, Thomas B., Kimberly M. Davis, Elena S. Lysenko, Alice Y. Zhou, Yimin Yu, and Jeffrey N. Weiser. 2010. ‘Recognition of Peptidoglycan from the Microbiota by Nod1 Enhances Systemic Innate Immunity’. Nature Medicine 16 (2): 228–31. 10.1038/nm.2087.

13. Crosswell, Lindsey C., and Janet M. Thornton. 2012. ‘ELIXIR: A Distributed Infrastructure for European Biological Data’. Trends in Biotechnology 30 (5): 241–42. 10.1016/j.tibtech.2012.02.002.

14. ECDC. 2020a. ‘AER EARS-Net Tables Showing Total Isolates Tested per Species’. ECDC. https://www.ecdc.europa.eu/sites/default/files/documents/EUEEA_tables_showing_total_isolates_tested_per_species.pdf.

15. ECDC. 2020b. ‘Antimicrobial Consumption in the EU - Annual Epidemiological Report 2019’. ECDC. https://www.ecdc.europa.eu/sites/default/files/documents/Antimicrobial-consumption-in-the-EU-Annual-Epidemiological-Report-2019.pdf.

16. Ero, Rya, Veerendra Kumar, Weixin Su, and Yong-Gui Gao. 2019. ‘Ribosome Protection by ABC-F Proteins—Molecular Mechanism and Potential Drug Design’. Protein Science : A Publication of the Protein Society 28 (4): 684–93. 10.1002/pro.3589.

17. Gacesa, R., A. Kurilshikov, A. Vich Vila, et al. 2022. ‘Environmental Factors Shaping the Gut Microbiome in a Dutch Population’. Nature 604 (7907): 732–39. 10.1038/s41586-022-04567-7.

18. Garay, Juan Antonio Raygoza, Williams Turpin, Sun-Ho Lee, et al. 2023. ‘Gut Microbiome Composition Is Associated With Future Onset of Crohn’s Disease in Healthy First-Degree Relatives’. Gastroenterology 165 (3): 670–81. 10.1053/j.gastro.2023.05.032.

19. Garcia-Gutierrez, Enriqueta, Melinda J. Mayer, Paul D. Cotter, and Arjan Narbad. 2019. ‘Gut Microbiota as a Source of Novel Antimicrobials’. Gut Microbes 10 (1): 1–21. 10.1080/19490976.2018.1455790.

20. Geese, Theresa, Corinna Bang, Andre Franke, Wolfgang Lieb, and Astrid Dempfle. 2025. ‘The Human Gut Microbiota in IBD, Characterizing Hubs, the Core Microbiota and Terminal Nodes: A Network-Based Approach’. BMC Microbiology 25 (1): 371. 10.1186/s12866-025-04106-0.

21. Ghosh, S, Mj Sadowsky, Roberts Mc, Gralnick Ja, and LaPara Tm. 2009. ‘Sphingobacterium Sp. Strain PM2-P1-29 Harbours a Functional Tet(X) Gene Encoding for the Degradation of Tetracycline’. Journal of Applied Microbiology 106 (4). 10.1111/j.1365-2672.2008.04101.x.

22. Goodrich, Julia K., Jillian L. Waters, Angela C. Poole, et al. 2014. ‘Human Genetics Shape the Gut Microbiome’. Cell 159 (4): 789–99. 10.1016/j.cell.2014.09.053.

23. Gradisteanu Pircalabioru, Gratiela, Mariana-Carmen Chifiriuc, Ariana Picu, Laura Madalina Petcu, Maria Trandafir, and Octavian Savu. 2022. ‘Snapshot into the Type-2-Diabetes-Associated Microbiome of a Romanian Cohort’. International Journal of Molecular Sciences 23 (23): 15023. 10.3390/ijms232315023.

24. Grigorescu, Andra, Florentina Dumitrescu, Stefania Dorobantu, et al. 2025. ‘An Epidemiological Survey of Sepsis in a Tertiary Academic Hospital from Southwestern Romania’. *Medicina (Kaunas*, Lithuania) 61 (4): 596. 10.3390/medicina61040596.

25. Heiman, Mark L., and Frank L. Greenway. 2016. ‘A Healthy Gastrointestinal Microbiome Is Dependent on Dietary Diversity’. Molecular Metabolism 5 (5): 317–20. 10.1016/j.molmet.2016.02.005.

26. Horn, Kadi J., Melissa A. Schopper, Zoe G. Drigot, and Sarah E. Clark. 2022. ‘Airway Prevotella Promote TLR2-Dependent Neutrophil Activation and Rapid Clearance of Streptococcus Pneumoniae from the Lung’. Nature Communications 13 (1): 3321. 10.1038/s41467-022-31074-0.

27. Horst, Rob ter, Martin Jaeger, Sanne P. Smeekens, et al. 2016. ‘Host and Environmental Factors Influencing Individual Human Cytokine Responses’. Cell 167 (4): 1111–1124.e13. 10.1016/j.cell.2016.10.018.

28. Hua, Xiaoting, Qian Liang, Min Deng, et al. 2021. ‘BacAnt: A Combination Annotation Server for Bacterial DNA Sequences to Identify Antibiotic Resistance Genes, Integrons, and Transposable Elements’. Frontiers in Microbiology 12 (July): 649969. 10.3389/fmicb.2021.649969.

29. Jeffery, Ian B., Denise B. Lynch, and Paul W. O’Toole. 2016. ‘Composition and Temporal Stability of the Gut Microbiota in Older Persons’. The ISME Journal 10 (1): 170–82. 10.1038/ismej.2015.88.

30. Krakau, Sabrina, Daniel Straub, Hadrien Gourlé, Gisela Gabernet, and Sven Nahnsen. 2022. ‘Nf-Core/Mag: A Best-Practice Pipeline for Metagenome Hybrid Assembly and Binning’. NAR Genomics and Bioinformatics 4 (1): lqac007. 10.1093/nargab/lqac007.

31. Lee, Kihyun, Sebastien Raguideau, Kimmo Sirén, et al. 2023. ‘Population-Level Impacts of Antibiotic Usage on the Human Gut Microbiome’. Nature Communications 14 (1): 1191. 10.1038/s41467-023-36633-7.

32. Lin, Qingqi, Yair Dorsett, Ali Mirza, et al. 2024. ‘Meta-Analysis Identifies Common Gut Microbiota Associated with Multiple Sclerosis’. Genome Medicine 16 (1): 94. 10.1186/s13073-024-01364-x.

33. Liu, Junfeng, Dongmin Hao, Xueyan Ding, et al. 2024. ‘Epidemiological Investigation and β-Lactam Antibiotic Resistance of Riemerella Anatipestifer Isolates with Waterfowl Origination in Anhui Province, China’. Poultry Science 103 (4): 103490. 10.1016/j.psj.2024.103490.

34. Lozupone, Catherine A., Jesse I. Stombaugh, Jeffrey I. Gordon, Janet K. Jansson, and Rob Knight. 2012. ‘Diversity, Stability and Resilience of the Human Gut Microbiota’. Nature 489 (7415): 220–30. 10.1038/nature11550.

35. Maeda, Yuichi, and Kiyoshi Takeda. 2017. ‘Role of Gut Microbiota in Rheumatoid Arthritis’. Journal of Clinical Medicine 6 (6): 60. 10.3390/jcm6060060.

36. McGinnis, Scott, and Thomas L. Madden. 2004. ‘BLAST: At the Core of a Powerful and Diverse Set of Sequence Analysis Tools’. Nucleic Acids Research 32 (Web Server issue): W20–25. 10.1093/nar/gkh435.

37. Mihele, Adina Ioana, Harrie Toms John, Nicoleta Negrut, Anca Ferician, Paula Marian, and Felicia Manole. 2025. ‘Hepatic Steatosis and Microbiota: A Regional Study on Patients from Western Romania’. Gastrointestinal Disorders 7 (1): 1. 10.3390/gidisord7010009.

38. Mirăuță, Bogdan, Cătălina Zenoaga-Barbăroșie, Monica Abrudan, et al. 2024. ‘From the Establishment of a National Bioinformatics Society to the Development of a National Bioinformatics Infrastructure’. 13:1002. Preprint, F1000Research, September 3. 10.12688/f1000research.153895.1.

39. Mulder, M., D. Radjabzadeh, J. C. Kiefte-de Jong, et al. 2020. ‘Long-Term Effects of Antimicrobial Drugs on the Composition of the Human Gut Microbiota’. Gut Microbes 12 (1): 1791677. 10.1080/19490976.2020.1791677.

40. Nurk, Sergey, Dmitry Meleshko, Anton Korobeynikov, and Pavel A. Pevzner. 2017. ‘metaSPAdes: A New Versatile Metagenomic Assembler’. Genome Research 27 (5): 824–34. 10.1101/gr.213959.116.

41. Parks, Donovan H., Maria Chuvochina, David W. Waite, et al. 2018. ‘A Standardized Bacterial Taxonomy Based on Genome Phylogeny Substantially Revises the Tree of Life’. Nature Biotechnology 36 (10): 996–1004. 10.1038/nbt.4229.

42. Popa, Laura Ioana, Irina Gheorghe, Ilda Czobor Barbu, et al. 2021. ‘Multidrug Resistant Klebsiella Pneumoniae ST101 Clone Survival Chain From Inpatients to Hospital Effluent After Chlorine Treatment’. Frontiers in Microbiology 11 (January). 10.3389/fmicb.2020.610296.

43. Rands, Chris M., Elizaveta V. Starikova, Harald Brüssow, Evgenia V. Kriventseva, Vadim M. Govorun, and Evgeny M. Zdobnov. 2018. ‘ACI-1 Beta-Lactamase Is Widespread across Human Gut Microbiomes in Negativicutes Due to Transposons Harboured by Tailed Prophages’. Environmental Microbiology 20 (6): 2288–300. 10.1111/1462-2920.14276.

44. Rodríguez-Molina, Daloha, Fanny Berglund, Hetty Blaak, et al. 2021. ‘Carriage of ESBL-Producing Enterobacterales in Wastewater Treatment Plant Workers and Surrounding Residents — the AWARE Study’. *European Journal of Clinical Microbiology & Infectious Diseases*, ahead of print, December 13. 10.1007/s10096-021-04387-z.

45. Ruiz-Limón, Patricia, Natalia Mena-Vázquez, Isabel Moreno-Indias, et al. 2022. ‘Collinsella Is Associated with Cumulative Inflammatory Burden in an Established Rheumatoid Arthritis Cohort’. Biomedicine & Pharmacotherapy 153 (September): 113518. 10.1016/j.biopha.2022.113518.

46. Schirmer, Melanie, Sanne P. Smeekens, Hera Vlamakis, et al. 2016. ‘Linking the Human Gut Microbiome to Inflammatory Cytokine Production Capacity’. Cell 167 (4): 1125–1136.e8. 10.1016/j.cell.2016.10.020.

47. Shoemaker, NB, H Vlamakis, K Hayes, and AA Salyers. 2001. ‘Evidence for Extensive Resistance Gene Transfer among Bacteroides Spp. and among Bacteroides and Other Genera in the Human Colon’. Applied and Environmental Microbiology 67 (2). 10.1128/AEM.67.2.561-568.2001.

48. Silva, Ana C. da, Jacob Lapkin, Qi Yin, Efrat Muller, and Alexandre Almeida. 2025. ‘Meta-Analysis of the Uncultured Gut Microbiome across 11,115 Global Metagenomes Reveals a New Candidate Biomarker of Health’. Preprint, bioRxiv, September 9. 10.1101/2025.09.09.675081.

49. Smith, Linda, Ekaterina Goldobina, Bianca Govi, and Andrey N. Shkoporov. 2023. ‘Bacteriophages of the Order Crassvirales: What Do We Currently Know about This Keystone Component of the Human Gut Virome?’ Biomolecules 13(4): 584. 10.3390/biom13040584.

50. Stamouli, Sofia, Moritz E. Beber, Tanja Normark, et al. 2023. ‘Nf-Core/Taxprofiler: Highly Parallelised and Flexible Pipeline for Metagenomic Taxonomic Classification and Profiling’. Preprint, bioRxiv, October 23. 10.1101/2023.10.20.563221.

51. Stojković, Vanja, María Fernanda Ulate, Fanny Hidalgo-Villeda, et al. 2019. ‘Cfr(B), Cfr(C), and a New Cfr-Like Gene, Cfr(E), in Clostridium Difficile Strains Recovered across Latin America’. Antimicrobial Agents and Chemotherapy 64 (1): e01074–19. 10.1128/AAC.01074-19.

52. Szekeres, Edina, Andreea Baricz, Cecilia Maria Chiriac, et al. 2017. ‘Abundance of Antibiotics, Antibiotic Resistance Genes and Bacterial Community Composition in Wastewater Effluents from Different Romanian Hospitals’. Environmental Pollution 225 (June): 304–15. 10.1016/j.envpol.2017.01.054.

53. Viprey, V. F., G. Granata, K. E. W. Vendrik, et al. 2023. ‘European Survey on the Current Surveillance Practices, Management Guidelines, Treatment Pathways and Heterogeneity of Testing of Clostridioides Difficile, 2018–2019: Results from The Combatting Bacterial Resistance in Europe CDI (COMBACTE-CDI)’. Journal of Hospital Infection 131: 213–20. 10.1016/j.jhin.2022.11.011.

54. Wood, Derrick E., Jennifer Lu, and Ben Langmead. 2019. ‘Improved Metagenomic Analysis with Kraken 2’. Genome Biology 20 (1): 257. 10.1186/s13059-019-1891-0.

55. Wu, Ziming, Mustasim Famous, Theano Stoikidou, et al. 2025. ‘Unravelling AMR Dynamics in the Rumenofaecobiome: Insights, Challenges and Implications for One Health’. International Journal of Antimicrobial Agents 66 (1): 107494. 10.1016/j.ijantimicag.2025.107494.

56. Yin, Qi, Ana C. da Silva, Francisco Zorrilla, Ana S. Almeida, Kiran R. Patil, and Alexandre Almeida. 2025. ‘Ecological Dynamics of Enterobacteriaceae in the Human Gut Microbiome across Global Populations’. Nature Microbiology 10 (2): 541–53. 10.1038/s41564-024-01912-6.

57. Yoshida, Chika, Mao Hagihara, Reina Azuma, et al. 2025. ‘Lipoproteins from Bilophila Wadsworthia Cell Wall Induce Innate Immune Responses through Toll-like Receptor 2’. Anaerobe 96 (September): 103001. 10.1016/j.anaerobe.2025.103001.

